# Amyloid ß Impacts Future Freezing of Gait in Parkinson’s Disease Via White Matter Hyperintensities

**DOI:** 10.1101/2020.10.29.360552

**Authors:** Mahsa Dadar, Janis Miyasaki, Simon Duchesne, Richard Camicioli

## Abstract

**Background:** Freezing of gait (FOG) is a common symptom in Parkinson’s Disease (PD) patients. Previous studies have reported relationships between FOG, substantia nigra (SN) degeneration, dopamine transporter (DAT) concentration, as well as amyloid β deposition. However, there is a paucity of research on the concurrent impact of white matter damage.

**Objectives:** To assess the inter-relationships between these different co-morbidities, their impact on future FOG and whether they act independently of each other.

**Methods:** We used baseline MRI and longitudinal gait data from the Parkinson’s Progression Markers Initiative (PPMI). We used deformation based morphometry (DBM) from T1-weighted MRI to measure SN atrophy, and segmentation of white matter hyperintensities (WMH) as a measure of WM pathological load. Putamen and caudate DAT levels from SPECT as well as cerebrospinal fluid (CSF) amyloid β were obtained directly from the PPMI. Following correlation analyses, we investigated whether WMH burden mediates the impact of amyloid β on future FOG.

**Results:** SN DBM, WMH load, putamen and caudate DAT activity and CSF amyloid β levels were significantly different between PD patients with and without future FOG (p < 0.008). Mediation analysis demonstrated an effect of CSF amyloid β levels on future FOG via WMH load, independent of SN atrophy and striatal DAT activity levels.

**Conclusions:** Amyloid β might impact future FOG in PD patients through an increase in WMH burden, in a pathway independent of Lewy body pathology.

## Introduction

Parkinson’s disease (PD) is a progressive neurodegenerative disorder primarily manifesting motor dysfunction, i.e. bradykinesia, tremor, muscular rigidity, impaired postural reflexes, and gait deficits ^1^. Freezing of gait (FOG) has been shown to develop in 25% of patients with early PD over two years of follow-up ^2^. FOG is characterized by a sudden transient and unexpected interruption in walking, usually during gait initiation or while turning ^3^. FOG increases the risk of falls and injury, and impacts the quality of life in PD patients, particularly in the later stages of the disease ^4^. At present, FOG poses an unmet need as neither medications nor advanced therapies such as deep brain stimulation reliably and significantly improve FOG outcomes. Understanding the determinants of future FOG might provide potential disease management and treatment opportunities.

Pathologically, PD leads to degeneration of the nigrostriatal dopaminergic pathway, substantial loss of substantia nigra (SN) neurons, and dopamine depletion ^1,5^. The hallmark of PD involves α-synuclein aggregations in the form of Lewy bodies initially in the SN region, leading to neuronal death ^1^. The degree of atrophy in the SN and presynaptic striatal dopamine depletion (as measured by DAT imaging) have been linked to FOG and gait impairment ^3,6–12^. Further, in patients with PD, amyloid beta (Aβ) deposition has been found to increase risk of FOG ^3,13,14^.

Of interest to our study, white matter hyperintensities (WMHs) are areas of increased signal on T2-weighted (T2w) magnetic resonance imaging (MRI) sequences, commonly observed in the aging population due to an interplay of ischemic, inflammatory, and protein deposition processes ^15,16^. In addition to cognitive impairment and executive dysfunction, WMHs have been associated with rigidity and gait impairment in individuals with ^7,17–23^ and without PD ^24–30^. Interestingly, amyloid β deposition has further been linked to an increase in the WMH burden through acceleration of processes such as neuroinflammation, reactive oxygen species production, and oxidative stress ^31–35^.

Given this inter-relationship between WMHs and amyloid β, and the fact that they both seem to impact gait in PD patients, a potential mechanism impacting gait in PD might involve increase in WMH loads due to amyloid β deposition. Since dopamine depletion and SN atrophy are more directly linked to Lewy body pathology, another factor to consider is whether WMH and amyloid β impact FOG through independently or through the dopamine/SN pathway.

In this study, we investigated the inter-relationships between different previously reported pathologic markers potentially affecting FOG in PD, namely SN atrophy, striatal DAT activity, amyloid β, and WMHs, and whether their impact on FOG is independent of each other or if there was mediation of effect on FOG. Specifically, we hypothesized that the impact of amyloid β on FOG might be mediated by WMH burden.

## METHODS

### Participants

We selected *de novo* PD patients from the Parkinson’s Progression Markers Initiative (PPMI) that had baseline T1-weighted (T1w) MRI and longitudinal clinical assessments available. PPMI is a longitudinal multi-center clinical study of *de novo* PD patients and age-matched healthy controls (http://www.ppmi-info.org) ^36^. The participants underwent comprehensive clinical and imaging assessments including Movement Disorders Society-Unified Parkinson’s Disease Rating Scale (MDS-UPDRS) ^37^.

### Ethics

The PPMI was approved by the institutional review board of all participating sites, and written informed consent was obtained from all participants included in the study.

### Freezing of Gait (FOG)

FOG was evaluated with MDS-UPDRS items 2.13 and 3.11. Future FOG was defined as whether the score for either item was ≥ 1 at any timepoint during the follow-up visits ^3^. MDS-UPDRS items for rest tremor (11 items: 2.10 and 3.15-18) and postural instability/gait difficulty (PIGD) designations (5 items: 2.12-13 and 3.10-12) were used to calculate future tremor and PIGD scores at 4-year follow-up ^38^. The tremor and PIGD scores were log-transformed to achieve a more normal distribution.

### Substantia Nigra Atrophy and WMH load measurements

#### MRI Preprocessing

All T1w images were preprocessed in three steps: noise reduction ^39^, intensity non-uniformity correction ^40^, and intensity normalization into range [0-100]. The preprocessed images were then linearly ^41^ registered to the MNI-ICBM152-2009c average template ^42^.

#### Substantia Nigra (SN) Atrophy

Deformation-based morphometry (DBM) was used as a marker of regional atrophy in the SN. DBM reflects macroscopic anatomical changes by spatially normalizing T1w MRIs to a standard space. To achieve this, the linearly registered T1w MRIs were non-linearly registered to the MNI-ICBM152-2009c template, composed of similar scans from 152 young, cognitively healthy individuals ^42^. The non-linear registration resulted in deformation fields sampled on a 1 mm^3^ grid ^43^. DBM maps were then calculated as the Jacobian determinant of the inverse deformation field ^44^. DBM values reflect the relative volume of a voxel to the corresponding voxel in the template; i.e. a value of 1 indicates similar volume to the same region in the template, values lower than one indicate volumes smaller than the corresponding region in the template, while values higher than one indicate volumes that are larger than the corresponding region in the template. Therefore, lower DBM values can be interpreted as indicative of lower structure volume, i.e. atrophy with respect to the (young) individuals of the template. The Xiao et al. deep grey matter atlas was used to calculate average DBM values in the SN at baseline ^45^.

#### WMH load measurement

Whole brain WMH load at baseline was used as a marker of WM pathology. WMHs were segmented based on the T1w images using a previously validated automatic segmentation technique that combines a set of location and intensity features obtained from a library of manually segmented scans with a Random Forest classifier to detect WMHs ^46–48^. The intensity features include voxel intensity, the probability of a specific intensity value being WMH or non-WMH, and the ratio of these two probabilities for the T1w images. Spatial features included a spatial WMH probability map indicating the probability of a voxel at a specific location being a WMH and the average intensity of a non-WMH voxel at that specific voxel location. The Random Forest classifier was then trained on a separate training library using these features to segment WMHs. The library used in this study was generated based on data from the Alzheimer’s Disease Neuroimaging Initiative (ADNI), since the PPMI T1w images had similar acquisition protocols. Once trained, the classifier was then applied to the study subject’s MRI. The WMH load used in this study was defined as the volume of the voxels identified as WMH in the standard space (in cm^3^) and are thus normalized for head size. WMH volumes were log-transformed to achieve normal distribution.

### MRI Quality Assessment

The quality of all the registrations and segmentations were visually assessed. All cases passed this quality control step. All MRI processing, segmentation and quality control steps were blinded to clinical outcomes.

### Dopamine Depletion

Dopamine transporter (DAT) imaging data with single-photon emission computed tomography (SPECT) was performed during the screening visit before enrollment in the PPMI study. We used the mean DAT uptake of the left and right caudate and putamen, as provided by the PPMI in table *DATScan_Analysis.csv,* as a marker of dopamine system integrity.

### CSF Amyloid Beta

CSF amyloid β_42_ measures at baseline were provided for PPMI participants with a lumbar puncture by the University of Pennsylvania. They used the the xMAP-Luminex platform with INNOBIAAlzBio3 immunoassay kit-based reagents (Fujirebio-Innogenetics, Ghent, Belgium) as markers of amyloid β deposition. The data are available in table *Current_Biospecimen_Analysis_Results.csv* provided by the PPMI. The CSF amyloid β values were also log-transformed to achieve normal distribution.

### Statistical Analyses

Partial correlations were used to assess the inter-relationships between different pathologic markers, controlling for age and sex. Regression models were used to investigate group differences between participants with and without future FOG, controlling for age and sex. Similarly, a secondary level regression analysis was performed to investigate the associations between the pathologic markers and future tremor and PIGD scores (4-year follow-up). A logistic regression model including all markers (i.e. SN DBM, putamen and caudate DAT levels, WMH load, and amyloid β) as well as age and sex was used to demonstrate the comparative impact of different measures on future FOG. From the evidence gathered between the correlations and the regression model, we used mediation analysis to test the hypothesis that WMH load mediates the effect of amyloid β on future FOG. More specifically, if WMH burden mediates much of the relationship between amyloid β levels and future FOG, we expected the relationship between amyloid β and future FOG to be reduced (partial mediation) or eliminated (full mediation) when the model accounts for the effect of WMH burden, implying a causal relationship between variables. To ensure that the impact of WMH and amyloid β on FOG is not through an inter-correlation with markers of Lewy body pathology, we controlled in the mediation analysis for putamen and caudate DAT levels and SN DBM. The logistic mediation analysis appropriate for binary outcomes was performed using MATLAB Mediation toolbox by Wager et al (https://github.com/canlab/MediationToolbox) ^49,50^. A 95% bootstrap confidence interval based on 10,000 bootstrap samples was used to estimate significance. All values were z-scored with respect to the population prior to regression and mediation analyses.

## RESULTS

Table 1 summarizes the baseline demographics and clinical characteristics of the participants enrolled in this study.

**Table 1.** Descriptive statistics for the participants enrolled in this study. Data are numbers (N) or mean ± standard deviation.

| Measure | Value |
| --- | --- |
| N | 423 |
| N<sub>Male</sub> | 274 |
| Age (years) | 61.80 ± 9.68 |
| MoCA | 27.10 ± 2.36 |
| MDS-UPDRSIII | 20.06 ± 9.47 |
| Future FOG (Yes/No) | 177/246 |
| CSF Amyloid β42 (pg/mL) | 915.6 ± 403.2 |
| SN DBM | 0.89 ± 0.09 |
| Putamen DAT (mean uptake) | 1.08 ± 0.70 |
| Caudate DAT (mean uptake) | 2.08 ± 0.74 |
| WMH Load (mm<sup>3</sup>) | 6.99 ± 1.07 |

MoCA= Montreal Cognitive Assessment. UPDRS= Unified Parkinson's Disease Rating Scale. FOG= Freezing Of Gait. CSF= CerebroSpinal Fluid. SN= Substantia Nigra. DBM= Deformation Based Morphometry. DAT= Dopamine Transporter. WMH= White Matter Hyperintensity.

### Inter-relationships Between Pathological Markers

Figure 1 shows the inter-correlations between different the pathological markers, controlling for age and sex. SN atrophy was significantly associated with putamen (r = 0.176, p = 0. 0004) and caudate (r = 0.222, p = 0. 0001) DAT levels, but not with CSF amyloid β. It was however significantly associated with WMH load (r = - 0.195, p = 0. 0001). As expected, DAT levels in putamen and caudate were highly correlated (r = 0.773, p < 0.0001), and were not associated with either CSF amyloid β levels or WMH loads. Finally, WMH loads were also significantly associated with CSF amyloid levels (r = - 0.199, p = 0. 0004).

**Figure 1.**
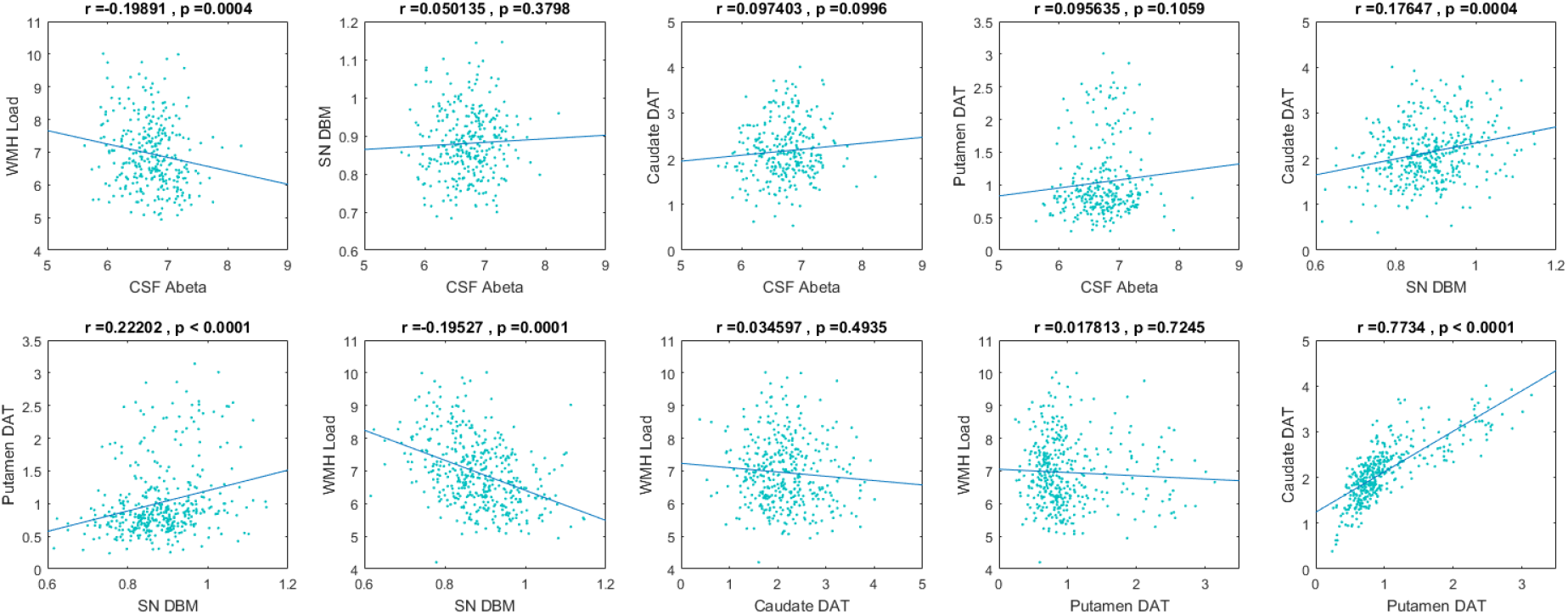
Intercorrelations between CSF amyloid, WMH load, substantia nigra atrophy, and putament and caudate DAT levels, correcting for age and sex. CSF= CerebroSpinal Fluid. Abeta= Amyloid β. SN= Substantia Nigra. DBM= Deformation Based Morphometry. DAT= Dopamine Transporter. WMH= White Matter Hyperintensity.

### Impact of Baseline Pathology on Future Freezing of Gait (FOG)

Figure 2 shows the differences in baseline biomarkers in patients with and without future FOG, controlling for age and sex. Patients with future FOG had significantly lower baseline SN DBM values (p = 0.003, indicating higher levels of atrophy), putamen (p = 0.002) and caudate (p = 0.0001) DAT levels, as well as significantly higher baseline WMH loads (p = 0.004), and significantly lower baseline CSF amyloid β levels (p = 0.008). The second level regression analysis showed significantly higher future tremor scores and lower future PIGD scores (p < 0.001) for patients with higher baseline putamen and caudate DAT levels, as well as significantly higher PIGD scores for patients with higher baseline WMH loads (p = 0.01). Future tremor scores were not associated with baseline WMH loads. Baseline SN DBM values and CSF amyloid β levels were also not significantly associated with future tremor or PIGD scores.

**Figure 2.**
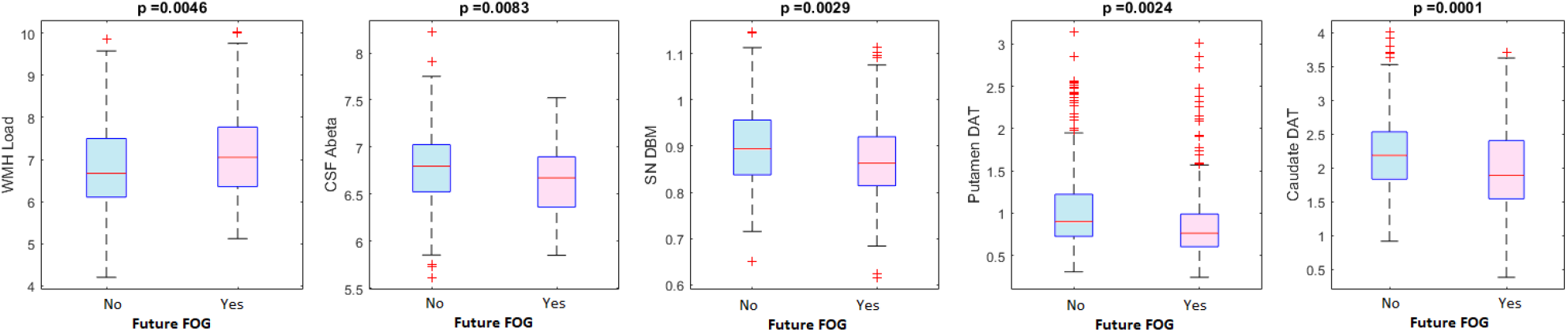
Differences in CSF amyloid, WMH load, substantia nigra atrophy, and putamen and caudate DAT levels between patients with and without future FOG, correcting for age and sex. FOG= Freezing Of Gait. CSF= CerebroSpinal Fluid. Abeta= Amyloid β. SN= Substantia Nigra. DBM= Deformation Based Morphometry. DAT= Dopamine Transporter. WMH= White Matter Hyperintensity.

Table 2 summarizes the results of the full logistic regression model including all the pathology markers, as well as age and sex. WMH load was the only measure that was significantly associated with future FOG (t = 2.495, p = 0.012), when including all the measures simultaneously in the model.

**Table 2.**
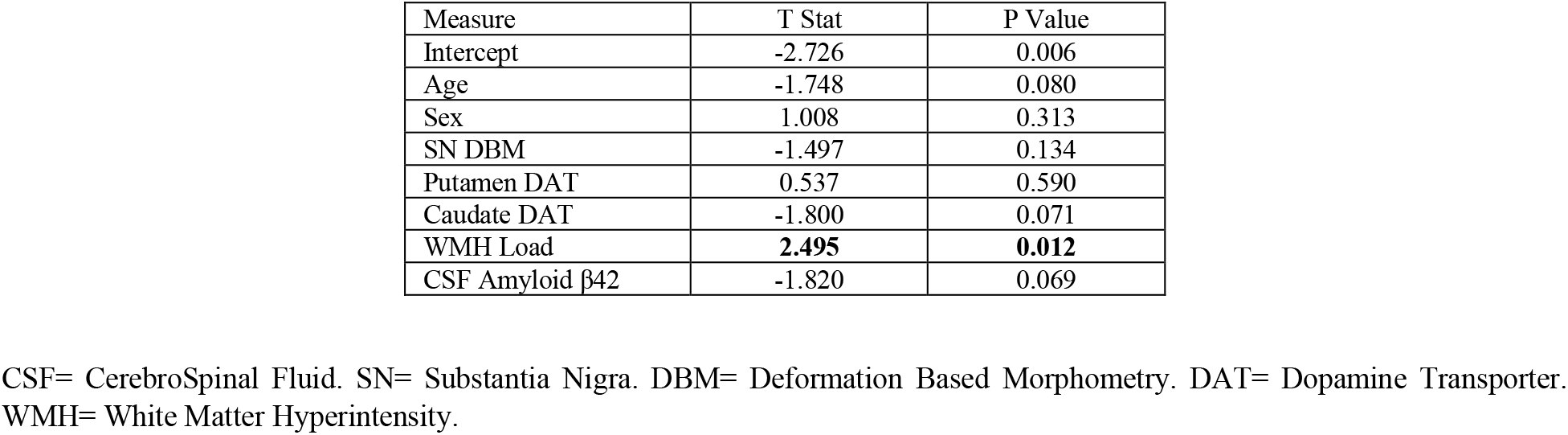
Logistic regression model results showing the impact of each pathology marker, as well as age and sex, on future freezing of gait.

### WMHs, Amyloid β, and FOG

Given (1) that amyloid β was only associated with WMH but not DAT levels or SN DBM, and (2) that only WMH was associated with future FOG in the regression model, we conducted a mediation analysis between between amyloid β, WMH load and future FOG (i.e. an effect of CSF amyloid levels on future FOG via WMHs), controlling for age, sex, SN DBM, putamen and caudate DAT levels. The analysis supported the existence of a full mediation (average causal mediation effect: ab = −0.013, 95%-CI: [−0.019, −0.009], p = 0.02; average direct effect (ADE): *C’* = −0.053, 95%-CI = [−0.071, −0.033], p = 0.053. Figure 3 summarizes the results of the mediation analysis, indicating CSF amyloid β impacts future FOG through an increase in WMH burden, independently of SN atrophy, striatal DAT activity, age, and sex.

**Figure 3.**
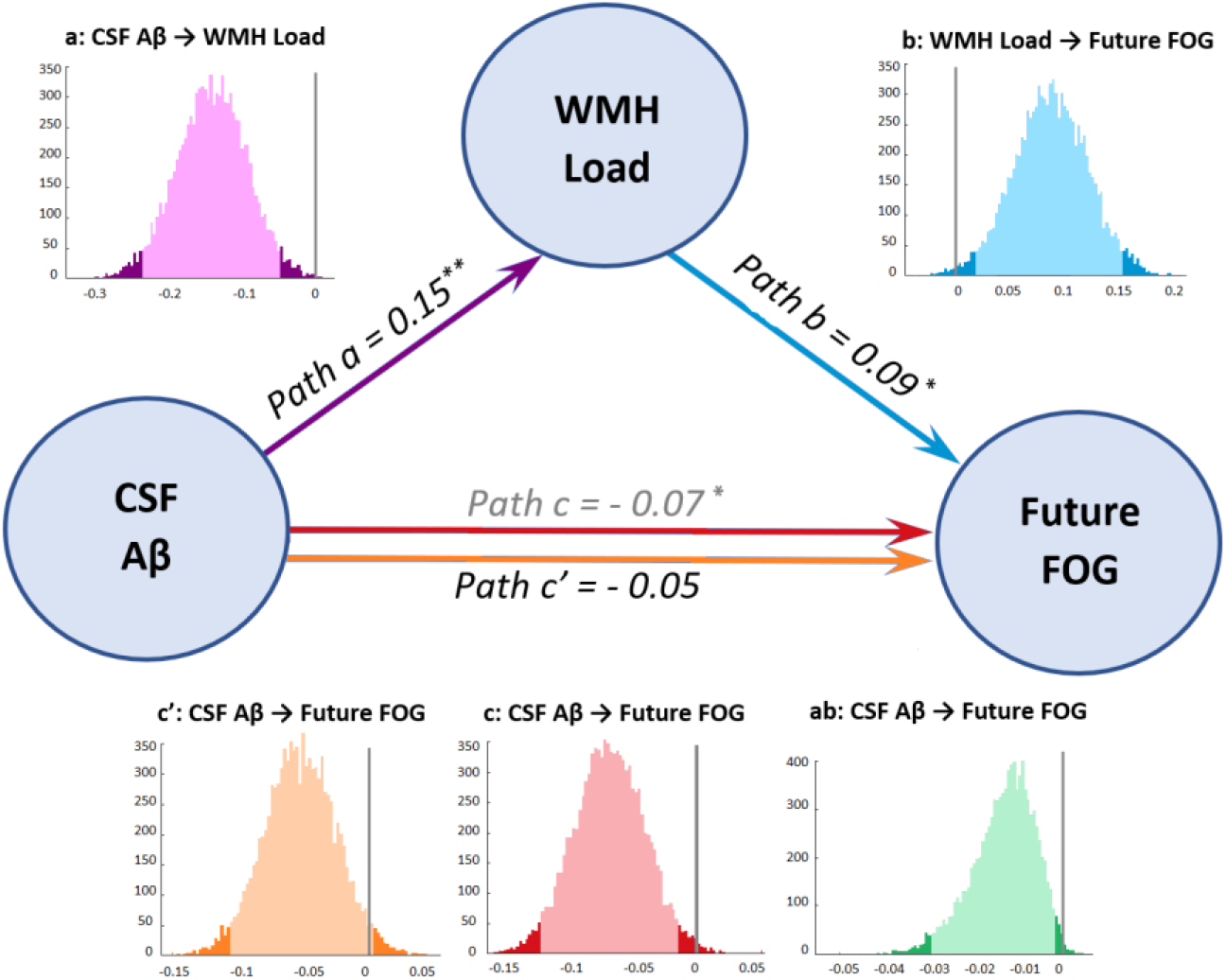
mediation analysis showing existence of a mediation effect of CSF amyloid β levels on future FOG via WMH load, correcting for age, sex, SN DBM, putamen and caudate DAT. CSF= CerebroSpinal Fluid. Abeta= Amyloid β, WMH= White Matter Hyperintensity.

## DISCUSSION

In this study, we investigated the impact of two previously known predictors of FOG, namely amyloid β and DAT levels, as well as two less well-established measures, WMH load and mean SN DBM, on future development of FOG in *de novo* PD patients from the PPMI dataset. When separately assessed, all variables were significantly different between PD patients that developed future FOG and those that did not. However, a follow-up mediation analysis showed that WMH load mediated the impact of amyloid β on future FOG, controlling for age, sex, DAT levels, and SN atrophy, indicating that amyloid β might impact future FOG through increasing WMH burden, in a pathway independent of Lewy body pathology.

DAT levels in putamen and caudate were highly correlated with each other and SN atrophy, but were not associated with either CSF amyloid β levels or WMH loads, indicating that amyloid β deposition and WMH progression might not be part of the Lewy body pathology in PD, and might impact gait through a separate pathway. A second level analysis investigating the relationship between the baseline pathology markers and future tremor and PIGD scores showed that WMH burden was associated with PIGD and not tremor, further supporting the hypothesis that WMHs have a distinct impact on axial symptoms, including PIGD.

The relationship between striatal DAT levels, amyloid β and FOG in PD patients has been previously shown, both in PPMI as well as other datasets ^3,6,8,9,13,14,51,52^. Regarding the SN, multiple recent studies have shown associations between free-water as an indirect measure of dopaminergic degeneration within the SN and higher Hoehn and Yahr stage, MDS-UPDRS III scores, and posture and gait in PD patients ^53–56^. Recently, in another longitudinal sample of PD patients in various stages of the disease, we reported that DBM is a sensitive marker of SN atrophy in PD and that SN DBM was associated with poorer motor performance in PD patients, in a separate cohort including PD patients with a full spectrum of illness ^7^. In the current sample, SN DBM was associated with future FOG, further demonstrating the value of assessing SN DBM as a T1w-based marker of PD pathology in future studies.

WMH burden is associated with gait deficits both in the aging population in general and in PD patients specifically ^7,17,20–22,51,52^. Given that WMH load is also associated with CSF amyloid β levels ^31,32,57,35^ and they both impact gait in PD, we hypothesized that they might impact future FOG through the same pathway. To investigate this, we used a mediation analysis, assessing the effect of WMH burden as the mediator on the relationship between amyloid β and future FOG. In addition to age and sex, we controlled for putamen and caudate DAT levels and SN DBM to ensure that the impact of WMH and amyloid β on FOG was not through an inter-correlation with markers of Lewy body pathology. Our results supported the existence of a full mediation, indicating that amyloid β impacts future FOG through an increase in WMH burden independent of SN atrophy and striatal DAT levels, suggesting that this might be an independent mechanism leading to gait impairment in PD.

The impact of WMHs on future FOG and its independence from striatal DAT activity is in line with another recent study by Chung et al. in a different cohort of de novo PD patients ^51^. Chung et al. showed that patients with moderate to severe WMH load (determined by visual rating) had a higher risk of developing FOG in the follow-up visits, adjusting for age, sex, and DAT activity ^51^. The present study confirms these findings in the PPMI dataset, using a more sensitive quantitative measure of WMH burden, and investigates the impact of amyloid β and SN atrophy in addition to WMHs and DAT activity. The mediation analysis results also suggest a potential mechanism for the WMHs impacting future FOG, through the same pathway as amyloid β, and independent of SN atrophy and DAT activity.

When including all the pathology markers as well as age and sex within the full logistic regression model, WMH load was the only signficiant predictor for future FOG, while the impact of caudate DAT levels and CSF amyloid β trended towards significance (P = 0.07). We included both putamen and caudate DAT levels in the model since they might represent changes in distinct circuits. However, since caudate and putamen DAT measures are highly correlated (r = 0.773, p < 0.0001), their effect on FOG might have been divided between the two variables, preventing either from reaching significance. To assess this, we performed the analysis only including either putamen or caudate DAT levels, and in this model, in addition to WMHs, caudate DAT levels (but not putamen) were also significantly associated with future FOG (t = - 2.15, p = 0.031), in line with previous studies ^9^. To ensure this inter-correlation does not impact the results of the mediation analysis, we also repeated the mediation analysis controlling for only either caudate or putamen DAT levels, and obtained similar results. Similarly, sex was not a significant predictor of future FOG in the full regression model, but when assessed alone (only correcting for age), males had a higher chance of future FOG than females (p = 0.03), in line with previous studies ^3^.

Since PIGD symptoms and FOG show partial response to dopaminergic medications ^58^, establishing a link between WMHs, PIGD and future FOG provides potential treatment considerations. Given that mobility is a strong influencer of quality of life and some of the risk factors for cerebrovascular disease and WMH progression are potentially modifiable through anti-hypertensive medications and lifestyle changes ^59,60^, PD patients with WMHs should have risk factors such as hypertension, hypercholesterolemia, and hyperglycemia treated in an effort to potentially mitigate gait impairment. However, a significant proportion of PD patients, particularly those with high cerebrovascular risk that may benefit from therapies such as statin, are still not prescribed such treatment ^61^. Taken together, our results suggest that development of FOG in *de novo* PD patients has a multi-pathway etiology, emphasizing the need for multi-modal therapeutic interventions in PD, including those targeting cerebrovascular risk factors.

## ACKNOWLEDGMENTS

MD is supported by a scholarship from the Canadian Consortium on Neurodegeneration in Aging in which SD and RC are co-investigators as well as an Alzheimer Society Research Program (ASRP) postdoctoral award. The Consortium is supported by a grant from the Canadian Institutes of Health Research with funding from several partners including the Alzheimer Society of Canada, Sanofi, and Women’s Brain Health Initiative. JMM is supported by the University Hospital Foundation, Dennis and Doreen Erker grant.

Data used in this article were obtained from the Parkinson’s Progression Markers Initiative (PPMI) database (www.ppmi-info.org/data). For up-to-date information on the study, visit www.ppmi-info.org. PPMI is sponsored and partially funded by the Michael J Fox Foundation for Parkinson’s Research and funding partners, including AbbVie, Avid Radiopharmaceuticals, Biogen, Bristol-Myers Squibb, Covance, GE Healthcare, Genentech, GlaxoSmithKline (GSK), Eli Lilly and Company, Lundbeck, Merck, Meso Scale Discovery (MSD), Pfizer, Piramal Imaging, Roche, Servier, and UCB (www.ppmi-info.org/fundingpartners). MD and DLC had full access to all the data in the study and takes responsibility for the integrity of the data and the accuracy of the data analysis.

## AUTHORS’ ROLES

Mahsa Dadar (PhD): Study concept and design, analysis and interpretation of the data and drafting and revising the manuscript.

Janis Miyasaki (MD): Interpretation of the data and revising the manuscript.

Simon Duchense (PhD): Study concept and design, interpretation of the data, drafting and revising the manuscript.

Richard Camicioli (MD): Study concept and design, interpretation of the data, drafting and revising the manuscript.

